# C2CAplus: a one-pot isothermal circle-to-circle DNA amplification system

**DOI:** 10.1101/2023.06.26.546530

**Authors:** Laura Grasemann, Paula Thiel Pizarro, Sebastian J. Maerkl

## Abstract

Rolling circle amplification (RCA) is a widely used DNA amplification method that uses circular template DNA as input and produces multimeric, linear single or double stranded DNA. Circle-to-circle amplification (C2CA) has further expanded this method by implementing product re-circularization using restriction and ligation, leading to a higher amplification yield, and enabling the generation of circular products. However, C2CA is a multistep, non-isothermal method, requiring multiple fluid manipulations and thereby compromises several advantages of RCA. Here, we improved C2CA to implement a one-pot, single step, isothermal reaction at temperatures ranging from 25 to 37°C. Our C2CAplus method is simple, robust, and produces large quantities of product DNA that can be seen with the naked eye.

## Introduction

RCA is a robust DNA amplification method that has become increasingly popular with a broad range of applications [1]. RCA requires a circular template. Sequence specific primers bind to this template and subsequently a polymerase with strand displacement activity, mainly bacteriophage phi29 DNA polymerase [2], will replicate the DNA in a rolling circle fashion, producing long multimeric strands of linear DNA. In contrast to PCR, no thermal cycling is required, which offers a multitude of advantages. DNA amplification using RCA is cost-effective, simple, and at the same time can be highly specific [3]. RCA has shown great potential for the use in diagnostics and as a biosensor, and various biologically relevant targets have been detected using RCA [4, 5, 6]. Hence, it is not surprising that isothermal systems such as RCA are anticipated to be used for on site testing in remote areas and poorly equipped healthcare settings [4]. Besides being used in biosensing, RCA is currently one of the most promising DNA replication methods for the construction of a synthetic cell [7, 8, 9, 10, 11]. To that end, however, a major drawback is that the output DNA structure of a RCA reaction is different to the input structure, that is linear instead of circular DNA.

Circle-to-circle-amplification (C2CA) was originally developed to improve RCA for the use in biosensing [12]. In a first step, padlock probes specific to the tested single-stranded DNA sequence and a ligase are used to generate a circular target DNA, which can then be amplified using standard RCA [13, 14]. In C2CA, the linear, single-stranded, multimeric RCA product is then digested by restriction digest and re-circularized in combination with a second primer by ligation generating circular monomeric products. These circles can then enter a consecutive round of RCA amplification, leading to circles of the opposite polarity, followed by another round of restriction, and ligation. The repetitive amplification, restriction, and ligation of template DNA during C2CA yielded 100x higher amounts of DNA than PCR [5]. Recent work has successfully demonstrated the detection of the Zika virus using padlock probes and C2CA combined with microfluidic affinity chromatography enrichment of the amplification product [15]. Martin and coworkers developed a biosensor to detect the antibiotic resistance gene sul1 for sulfonamide resistance using padlock probes. They furthermore developed a readout method that was visible by the naked eye using functionalized magnetic nanoparticles that aggregate with the C2CA product [16].

While RCA has the advantage that the process is simple, isothermal and is functional at ambient temperatures, current C2CA methods lost these advantages, as different temperatures are required for restriction and ligation, additional heat inactivation steps are necessary, and multiple reagent additions are needed in each round of amplification. C2CA therefore requires considerable user interaction or the use of liquid handling robotic platforms. We thus set out to develop an isothermal C2CA system that contains all components in a single tube, creating an isothermal C2CA method that requires no user interaction or complex automation. Our C2CAplus method is more sensitive than standard RCA, and produces large quantities of DNA that can be seen by naked eye. C2CAplus functions robustly in a temperature range between 25-37°C and produces circular DNA products that can be transformed. We anticipate that C2CAplus will pave the way towards lower-cost, and easier-to-use RCA based biosensing methods, and could form the basis of a DNA replication system in a synthetic cell.

## Results

### Isothermal restriction and ligation

In RCA one or more primers bind to a circular target DNA. A strand displacing DNA polymerase elongates these primers in a rolling circle manner producing long, multimeric, initially single-stranded DNA products. If an anti-directional primer is added, double-stranded, multimeric DNA can be generated. In C2CA and C2CAplus a restriction enzyme is then used to monomerize this double-stranded DNA, and subsequently a ligase ligates the ends, thus re-circularizing the initially linear product (Figure 1A).

**Figure 1.**
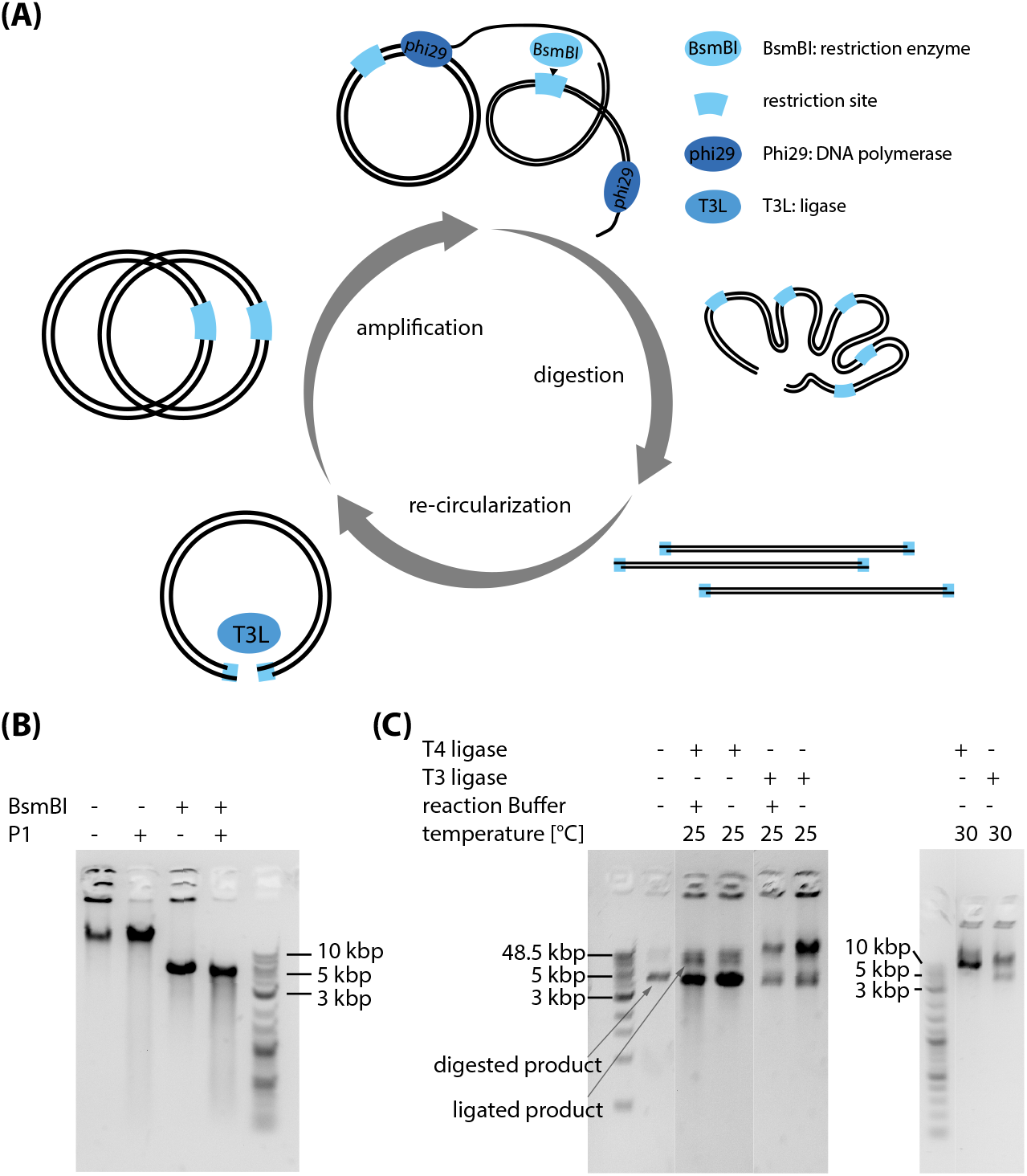
: **(A)** Schematic of a C2CAplus amplification scheme that integrates DNA amplification, restriction, and re-circularization. DNA amplification by Phi29 DNA polymerase produces multimeric doublestranded DNA that can be monomerized using a restriction enzyme. These monomers are subsequently ligated and thus circularized using a ligase. **(B)** Characterization of restriction digestion of an amplification product. The left band shows the untreated reaction with product in the pocket and a distinct band at a running length greater than the uppermost band of 48.5 kbp of the 1kb Extend Ladder. The product in the pocket can be digested by exonuclease P1, which is specific for ssDNA. The band at *>* 48.5 kbp can be cut by BsmBI to a length of around 5 kbp, which is the monomeric length of our input plasmid, indicating that this band is multimeric dsDNA. **(C)** Ligation activity of T4 and T3 DNA ligases after an initial BsmBI digest with and without reaction buffer, at 25°C and 30°C. A non-cropped version of the left gel is shown in Supplementary Figure S2.

To develop an isothermal one-pot C2CAplus system, one thus requires a restriction enzyme and ligase that operate under isothermal conditions, ideally around 30°C. We used the enzyme BsmBI v2 (NEB) to cut the sequence of our target plasmid exactly once to monomerize the double-stranded product, as can be seen by the single band in Figure 1B. The non-digested high-molecular weight product in the gel loading pocket is most likely single-stranded DNA that can’t be cut by BsmBI, but can be digested by P1, a nuclease that specifically degrades ssDNA and RNA (Figure 1B). Adding both P1 and BsmBI produced a single band at around 5 kbp (Figure 1B), which is presumably monomeric, double-stranded DNA. Albeit optimal conditions for BsmBI v2 include NEB buffer 3.1 and incubation at 55°C, BsmBI activity at 30°C in an RCA reaction environment is sufficient to monomerize all product DNA after 18 h of incubation at 30°C using a concentration of around 0.5 -0.2 U/μL (Supplementary Figure S1).

In the next step we digested an RCA product using BsmBI and screened several commercially available ligases including T4 DNA ligase, Taq DNA ligase, T3 DNA ligase, T7 DNA ligase, and *E. coli* DNA ligase, under their respective optimal conditions. When transforming the ligation product into 10-beta competent *E. coli* cells, we observed a significant increase in colonies for T4 DNA ligase, T3 DNA ligase, and *E. coli* DNA ligase treated samples compared to the BsmBI digested control and the plain RCA product (Supplementary Figure S3). As the optimal temperatures for T3 and T4 ligases of 25°C are already close to our target temperature of 30°C, and *E. coli* DNA ligase optimally operates at 16°C, we focused on T3 and T4 ligase.

Optimal ligation conditions for both T3 and T4 ligases include the above mentioned ligation temperature of 25°C, addition of a reaction buffer, and dilution of the restricted RCA product. We set out to investigate the performance of both ligases under non-optimal conditions. We added the ligases to the non-diluted restricted RCA product and omitted the respective reaction buffer. Both ligases retained activity under these conditions (Figure 1C). In a second experiment we increased the ligation temperature from the optimal 25°C to 30°C. Both, T3 and T4 DNA ligases ligated the restricted product at 30°C in absence of ligase buffer at both temperatures, rendering them promising candidates for an isothermal one-pot C2CA system.

### One-Pot C2CAplus

The development of a one-pot C2CAplus system requires a careful balance of restriction enzyme to ligase ratio (Figure 2A). If too little restriction enzyme is present, most product will remain multimeric, rendering re-circularization inefficient. If too much restriction enzyme is present, the restriction enzyme will linearize all DNA including newly ligated circular DNA. As RCA only works on circular DNA, the presence of only linear template will not lead to an improved DNA amplification. We thus tested concentrations between 0 -0.15 U/μL of BsmBI which were below the 0.2 U/μL used above and which digested all RCA product during the time course of a RCA reaction. As T3 out-performed T4 ligase in our initial characterization (Figure 1C), we chose to test different concentrations of T3 DNA ligase between 0-120 U/μL. To develop our one-pot C2CAplus reaction, we investigated a combinatoric space of BsmBI-to-T3 concentrations (Figure 2B).

**Figure 2:**
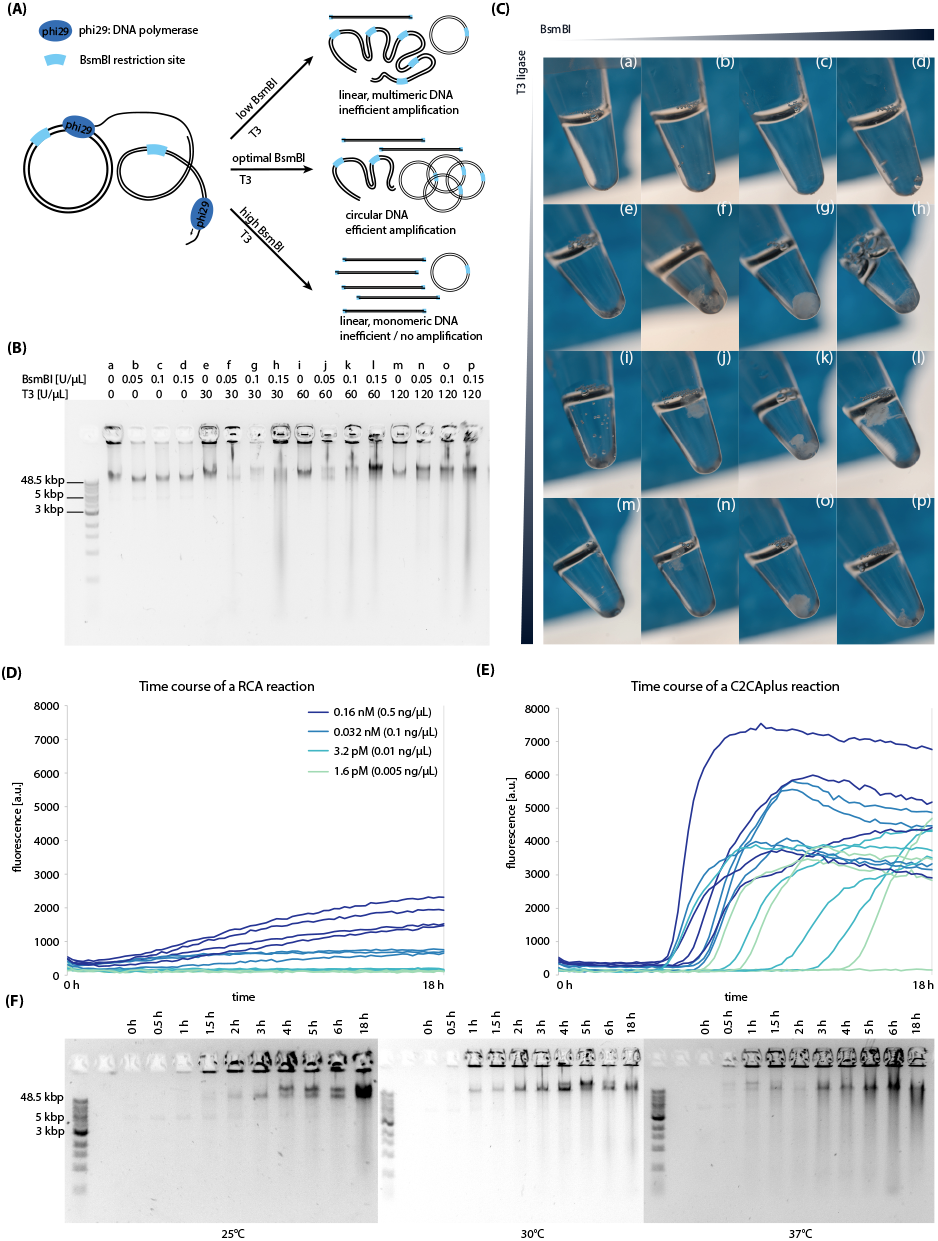
**(A)** Schematic description of possible outcomes of C2CAplus reactions. If the BsmBI concentration is too low, most of the RCA product will remain multimeric leading to inefficient amplification. If the BsmBI concentration is too high, the restriction enzyme will monomerize the majority of the product and input DNA, which will reduce amplification. Similarly, optimal concentrations of ligase are required to facilitate efficient re-circularization of the digested products present in the reaction, but is not as critical as the restriction enzyme concentration. If the BsmBI-to-ligase ratio is optimal, mostly circular product will be generated, which serves as an efficient template for further rounds of replication, thus generating a high amplification rate. **(B)** Agarose gel showing the products of C2CAplus reactions with different concentrations of BsmBI and T3 DNA ligase **(C)** Pictures of the reaction tubes from (B). **(D), (E)** RCA (D) and C2CA (E) reaction curves measured on a platereader using different input plasmid concentrations. C2CAplus successfully amplifies lower DNA input concentration than RCA. **(F)** Agarose gels of a time course of C2CAplus using different reaction temperatures.

The tested BsmBI concentrations were indeed low enough to not drastically affect amplification (Figure 2B). However, without the addition of T3 ligase, amplification efficiency was decreased compared to the control lacking T3 ligase and BsmBI (Figure 2B, first band). The sole addition of T3 ligase did not change amplification efficiency considerably (Figure 2B, bands e, i, m). However, if both BsmBI and T3 ligase were added together, amplification efficiency was increased (Figure 2B, bands f-h, j-l, n-p).

Amplification was generally so efficient that the solution in the reaction tube became highly viscous and was difficult to be applied to an agarose gel. Amplification was so large with optimal combinations of BsmBI and T3 ligase that a white, highly viscous precipitate formed in the reaction tubes, which became more pronounced after a freeze -thaw cycle (Figure 2C). Precipitate formation could be seen for all reactions containing both BsmBI and T3 ligase, indicating that C2CAplus is robust over the tested range of restriction enzyme-to-ligase ratios. However, a concentration of 0.1 U/μL BsmBI with T3 ligase at 30, 60, and 120 U/μL, seemed to be especially advantageous for precipitate formation, with a qualitative optimum at 0.1 U/μL BsmBI and 30 U/μL T3 ligase (Figure 2C panels g, k, o). Precipitate formation is less pronounced at lower and higher BsmBI concentrations, indicating that BsmBI concentration is critical for DNA amplification efficiency. It should be mentioned that although the precipitate can be seen directly after the reaction, precipitate becomes even more prominent after freeze-thawing the reaction tubes. We subsequently validated DNA sequence integrity by transforming 10-beta *E. coli* cells with the C2CAplus product, isolated the plasmid via Miniprep, and sequence verified the plasmid.

### C2CAplus is highly sensitive and robust

We tested the sensitivity of C2CAplus by titrating input plasmid concentrations between 1.6 pM and 0.16 nM (0.005 ng/μL and 0.5 ng/μL) to an RCA and a C2CAplus reaction supplemented with EvaGreen DNA stain and measured fluorescence over time (Figure 2D). While plain RCA produced an increase in fluorescence for input plasmid concentrations of 0.5 and 0.1 ng/μL (Figure2D), C2CAplus amplified plasmid DNA for all tested input DNA concentrations including 0.005 ng/μL (Figure 2E). C2CAplus also generated a significantly higher fluorescent signal compared to plain RCA. The curves for C2CAplus were exponential, in contrast to the linear curves obtained for plain RCA, indicating that the restriction enzyme and ligase indeed circularize a significant amount of the RCA product, which can subsequently serve as a circular input template for further rounds of replication leading to an exponential amplification.

If used as a biosensor or in a synthetic cell, a tolerance to reaction temperatures is ideal. We therefore compared the performance of C2CAplus at 30°C to its performance at 25°C and 37°C. Figure 2F shows the time course of C2CAplus reactions at 25°C, 30°C, and 37°C. Despite the reaction at 25°C being delayed by about 30 min compared to the reactions at higher temperatures, no differences are observed between the general amplification levels, indicating that C2CAplus performs robustly in the tested temperature range of 25 to 37°C.

## Discussion

In this work we developed a fully isothermal one-pot C2CA method called C2CAplus using the restriction enzyme BsmBI to cut the linear multimeric DNA produced by phi29 DNA polymerase, and T3 DNA ligase to re-ligate the monomers to form circular DNA. As the only sequence requirement for C2CAplus is a single unique restriction site, the method promises to be versatile and broadly applicable for various purposes. C2CAplus combines the main advantage of standard RCA by being able to operate in isothermal conditions, with the advantages of C2CA of a higher amplification rate and the ability to generate circular DNA products.

C2CAplus successfully amplified DNA with input concentrations of 0.005 ng/μL (1.6pM). The exponential amplification produces large quantities of DNA and generates precipitate that is visible to the naked eye. We anticipate that a simple readout based on precipitate formation could facilitate the use of C2CAplus as a biosensor in remote areas, rendering readout independent of sophisticated laboratory equipment. However, comparable to LAMP and other highly sensitive DNA amplification methods that exponentially amplify DNA, C2CAplus is also prone to producing false-positives and care must be taken to avoid DNA contamination when used as a biosensor, or highly specific primers need to be developed.

DNA hydrogels have become popular due to characteristics that render them advantageous especially in biomedical applications and biosensing [17]. Recently, Song and coworkers demonstrated the fabrication of a DNA hydrogel in 24 h, using multiple primed RCA reactions [18]. As our C2CAplus method produces large quantities of DNA that precipitates in the tube to a white, viscous, gel-like structure, we expect that C2CAplus could also be helpful in efficiently producing DNA for use in biomaterials.

Lastly, we see potential for C2CAplus to be used as a DNA replication mechanism in a synthetic cell. Thus far, endeavours have mainly focused on implementing standard RCA in cell-free transcription/translation systems [7, 9, 10] and Van Nies and coworker implemented a method that replicates DNA from a linear template [19]. One main challenge of using RCA as a replication method is the structural difference of DNA produced by RCA: while the input DNA structure is circular, the output structure is linear and multimeric. Sakatani and coworkers [8], as well as Okauchi and coworkers [11] attempted to solve this issue by implementing a re-circularization mechanism using cre recombinase. However, cre recombinase seemed to inhibit DNA replication. Okauchi [11] solved this by evolving the DNA. Yet, in the larger context of eventually building an artificial cell, sequence restrictions to avoid inhibition by a cre recombinase may be problematic. We therefore suggest that implementing re-circularization by restriction and re-ligation as introduced here may be a viable path towards achieving integrated DNA replication in a cell-free transcription -translation system.

## Methods

### Stand-alone RCA, restriction, and ligation reactions

A standard RCA reaction was performed using 0.1 U/μL phi29 DNA polymerase (ThermoFisher Scientific, stock: 10 U/μL) supplemented with 1x reaction buffer, 1 μM 3’ final primer and 1 μM 5’final primer, (sequences in Supplementary Table 1), 0.5 mM dNTP mix (ThermoFisher Scientific), and an input plasmid (sequence in Supplementary Information) concentration of 0.5 ng/μL (0.16 nM) unless indicated otherwise.

Nuclease P1 (NEB) treatment was performed as recommended by the supplier in a reaction volume of 10 μL, by supplementing 8.5 μL of RCA product with 1 μL NEB 1.1 reaction buffer (final concentration: 1x) and 0.5 μL P1 nuclease (final concentration: 5 U/μL). Restriction digest using BsmBI v2 (NEB, stock concentration 10 000 U/mL) was performed at a BsmBI-v2 concentration of 0.67 U/μL. The restriction was performed in undiluted RCA product omitting any additional buffer at 55°C for 1 h, followed by heat inactivation at 80°C, unless indicated otherwise.

All ligases (T3, T4, *E*.*coli*, T7, Taq) used during this work were obtained from NEB. Ligations were performed as indicated by the supplier. For the transformation experiments in Supplementary Figure S3, 1 μL of BsmBI digested RCA product was supplemented with 0.5 μL of each ligase in 1x reaction buffer in a total reaction volume of 10 μL. Incubation times and temperatures are as follows: T4 ligase was incubated at 25°C for 2 h, followed by heat inactivation at 65°C for 10 min; T3 and T7 ligases were incubated at 25°C for 30 min without heat inactivation; Taq ligase was incubated at 45°C for 15 min; *E. coli* ligase was incubated at 16°C for 30 min, followed by heat inactivation at 65°C for 20 min.

For transformation reactions, 5 μL of the respective ligated product was added to one vial (50 μL) of NEB 10-beta competent *E. coli* cells. Non-ligated RCA and BsmBI products were diluted 1:10 in MiliQ water prior to transformation, to obtain the same dilution factor as the ligated product. For the ligation tests of T3 and T4 ligases in Figure 1C, 4.5 μL of BsmBI digested RCA product was added to the reactions containing reaction buffer, and 9.5 μL of BsmBI digested RCA product was added to the reactions without reaction buffer. Reaction temperatures and times are as indicated above, unless specified otherwise.

Gel electrophoresis was performed using 1 % agarose gels and the ladders used were either Quick-Load 1 kb Extend DNA Ladder (NEB), or Quick-Load Purple 1 kb plus DNA Ladder (NEB).

### C2CAplus reactions and platereader experiments

C2CAplus reactions were performed in a total reaction volume of 20 μL or 50 μL. 0.1 U/μL phi29 DNA polymerase was supplemented with 1x reaction buffer, 0.5 mM dNTP mix, 1 μM 3’final primer, and 1 μM 5’final primer (Supplementary Table 1). The reaction was additionally supplemented with 120 U/μL T3 ligase, and 0.1 U/μL BsmBI, unless indicated otherwise. It should be noted that T3 DNA ligase is ATP dependent. No ATP is present in the RCA reaction buffer, and by omitting the T3 ligase buffer, we do not add any ATP to the reaction. We suppose that contaminating ATP fuels the ligation reaction. The C2CAplus reaction was incubated in a thermocycler for 18 h at 30°C. The highly viscous product was subsequently applied to a 1% agarose gel for analysis. It needs to be noted that due to the high viscosity of the product, the application to a gel was challenging. For the analysis of precipitate formation in Figure 2B, the product was freeze-thawed once before imaging. Here, the total reaction volume was 50 μL. All platereader experiments were performed with 20 μL volumes and the reactions were further supplemented with 2% BSA and 0.25x EvaGreen (Biotium, stock: 20x in H2O) in an optical, black, flat bottom 384 well-plate (Thermo Scientific). It should be noted that EvaGreen inhibits the reaction if applied at higher concentrations, and the addition of BSA is crucial in platereader experiments. Samples were sealed using a SealPlate film (Excel Scientific). Fluorescence was measured every 10 min at 500 / 530 nm ex/em for 18 h at 30°C, and during every 10 min interval, the plate was shaken for 10 s.

### Image and data processing

Gel images were cropped using Adobe Photoshop and labeled in Adobe Illustrator. Plate images in Figure S3 were adjusted in brightness and contrast for better visibility of the colonies using Fiji.

## Author Contributions

L.G. and P.T. performed experiments. L.G., P.T., and S.J.M. designed experiments. L.G. and S.J.M analyzed data, and wrote the manuscript.

## Acknowledgements

The authors would like to express their gratitude to Barbora Lavickova for helpful input and comments, and to Fabien Jammes for his help with the pictures of the DNA precipitate.

## Supplementary Information

**Figure S1:**
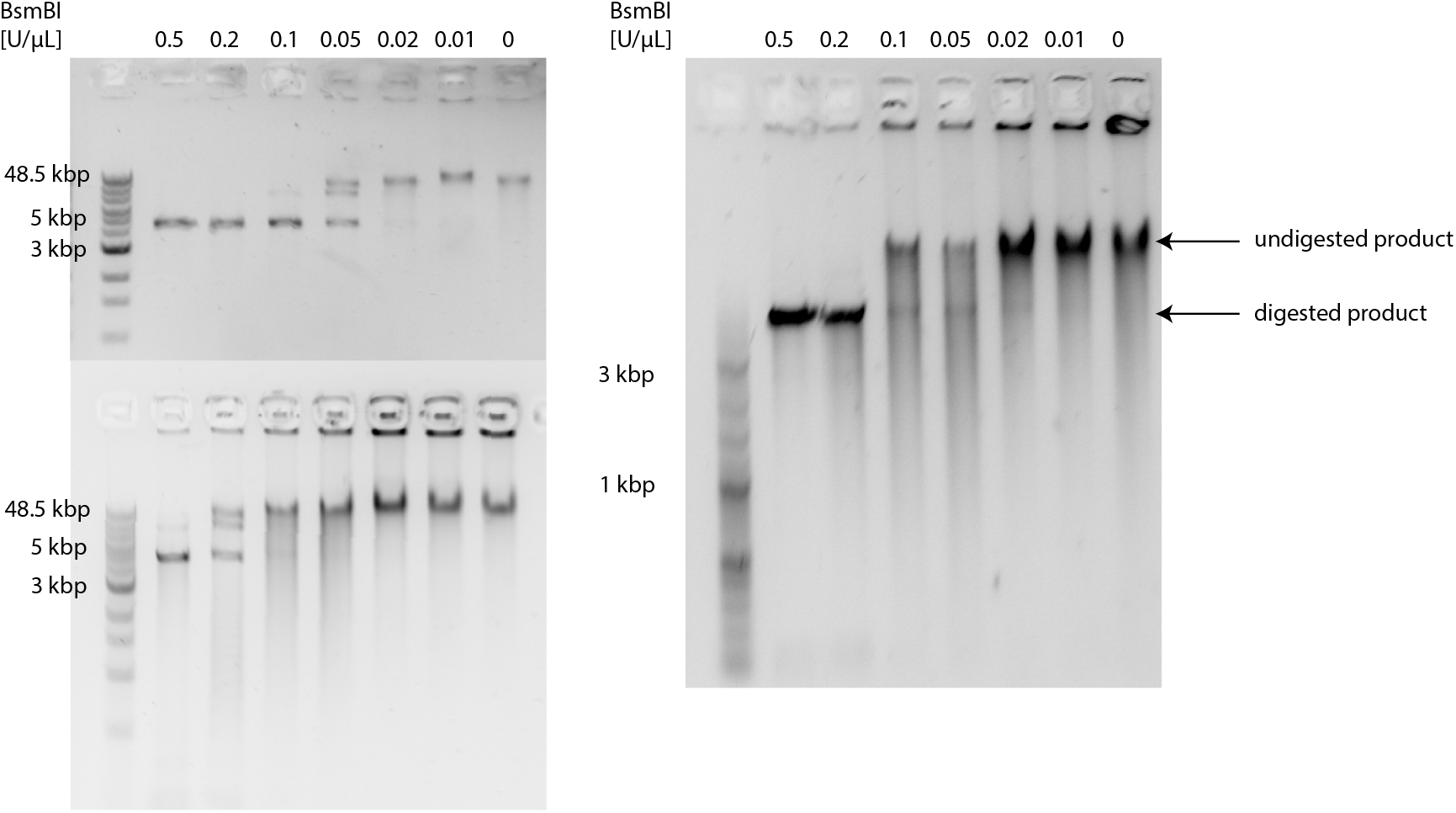
BsmBI titration at 30°C and 18 h reaction time. Three independent replicates were performed. A concentration of 0.5 to 0.2 U/μL digests most of the RCA product in the given conditions.

**Figure S2:**
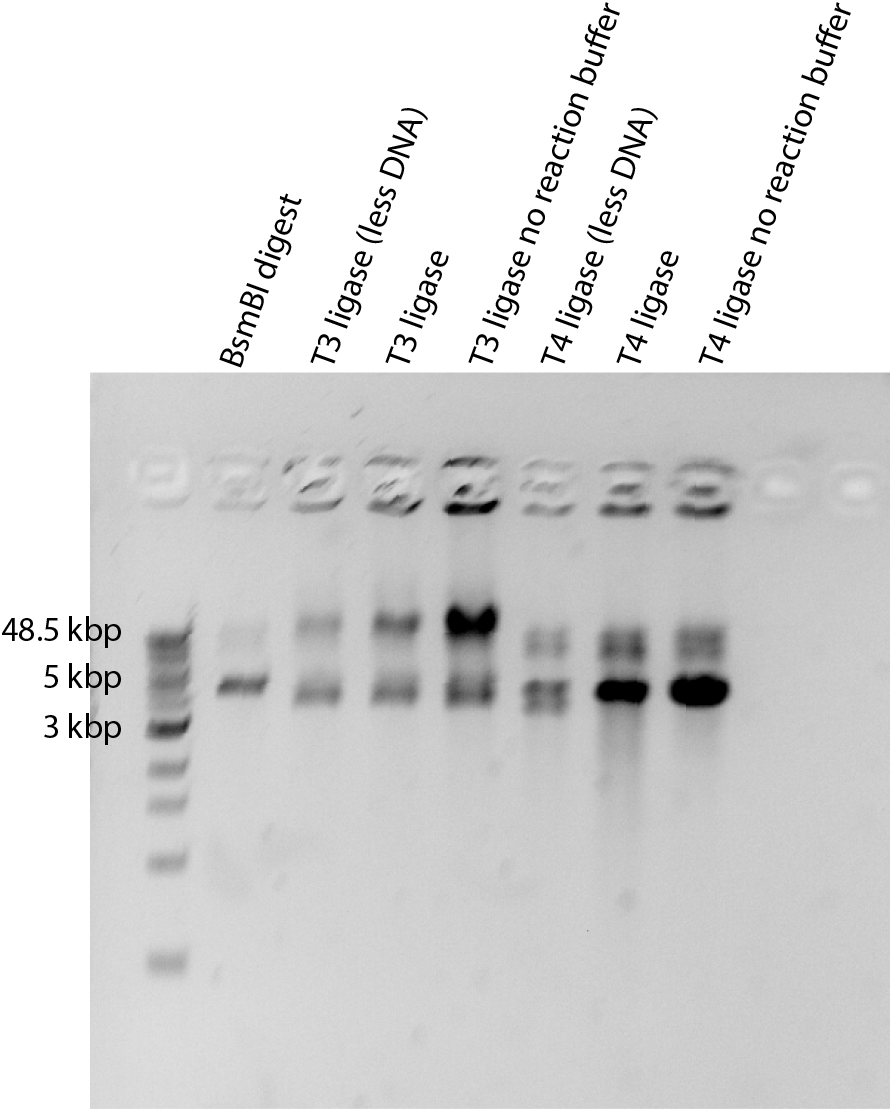
Non-cropped gel of Figure 1C. The first lane is a BsmBI digested product as shown in Figure 1C. The second lane is a T3 ligase ligation reaction with less DNA (3 μL in a total volume of 10 μL), not included in Figure1C. The following two lanes are again included in Figure 1C, and show the T3 ligation with and without reaction buffer. The 5th lane, T4 ligase (less DNA) is not included in Figure 1C, and shows the T4 ligation with 3 μL of DNA in a total volume of 10 μL. The last two lanes are again included in Figure 1C, and show the T4 ligation reaction with and without reaction buffer.

**Figure S3:**
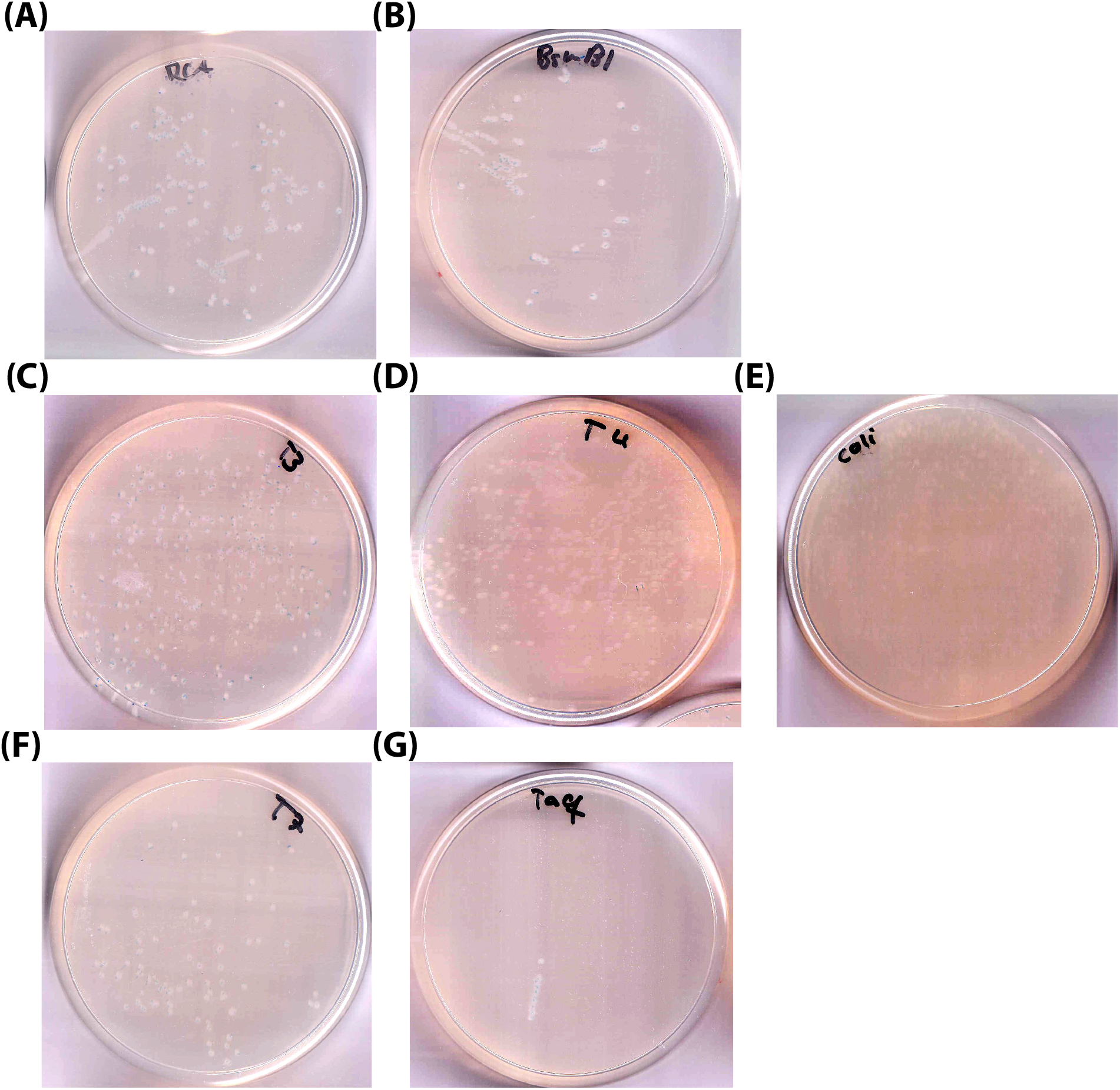
Ligase screen. Transformation results of: (A) non-treated RCA product, (B) BsmBI digested RCA product, T3 (C) T3 ligase, T4 (D) T4 ligase, (E) *E. coli* ligase, (F) T7 ligase, (G) Taq ligase.

### DNA sequences

#### Primer sequences

The primers used in this work were:

**Table 1:**
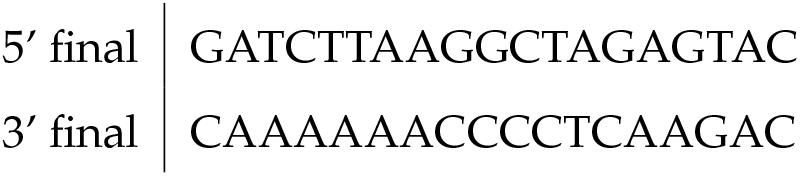
Primers used during this work

#### Plasmid sequence

The plasmid used in this work is a pSTBlue plasmid encoding Kanamycin and Ampicillin resistance as well as an EGFP gene. The BsmBI restriction site is located inside the Kanamycin resistance gene. The complete plasmid sequence is:

CTCAGGCGCAATCACGAATGAATAACGGTTTGGTTGATGCGAGTGATTTTGATGACGAGCGTAATGGCTGGCCTGTTGAACAAGTCTGGAAAGAAATGCATAAACTTTTGCCATTCTCACCGGATTCAGTCGTCACTCATGGTGATTTCTCACTTGATAACCTTATTTTTGACGAGGGGAAATTAATAGGTTGTATTGATGTTGGACGAGTCGGAATCGCAGACCGATACCAGGATCTTGCCATCCTATGGAACTGCCTCGGTGAGTTTTCTCCTTCATTACAGAAACGGCTTTTTCAAAAATATGGTATTGATAATCCTGATATGAATAAATTGCAGTTTCATTTGATGCTCGATGAGTTTTTCTAAGAATTAATTCATGACCAAAATCCCTTAACGTGAGTTTTCGTTCCACTGAGCGTCAGACCCCGTAGAAAAGATCAAAGGATCTTCTTGAGATCCTTTTTTTCTGCGCGTAATCTGCTGCTTGCAAACAAAAAAACCACCGCTACCAGCGGTGGTTTGTTTGCCGGATCAAGAGCTACCAACTCTTTTTCCGAAGGTAACTGGCTTCAGCAGAGCGCAGATACCAAATACTGTCCTTCTAGTGTAGCCGTAGTTAGGCCACCACTTCAAGAACTCTGTAGCACCGCCTACATACCTCGCTCTGCTAATCCTGTTACCAGTGGCTGCTGCCAGTGGCGATAAGTCGTGTCTTACCGGGTTGGACTCAAGACGATAGTTACCGGATAAGGCGCAGCGGTCGGGCTGAACGGGGGGTTCGTGCACACAGCCCAGCTTGGAGCGAACGACCTACACCGAACTGAGATACCTACAGCGTGAGCTATGAGAAAGCGCCACGCTTCCCGAAGGGAGAAAGGCGGACAGGTATCCGGTAAGCGGCAGGGTCGGAACAGGAGAGCGCACGAGGGAGCTTCCAGGGGGAAACGCCTGGTATCTTTATAGTCCTGTCGGGTTTCGCCACCTCTGACTTGAGCGTCGATTTTTGTGATGCTCGTCAGGGGGGCGGAGCCTATGGAAAAACGCCAGCAACGCGGCCTTTTTACGGTTCCTGGCCTTTTGCTGGCCTTTTGCTCACATGTTCTTTCCTGCGTTATCCCCTGATTCTGTGGATAACCGTATTACCGCCTTTGAGTGAGCTGATACCGCTCGCCGCAGCCGAACGACCGAGCGCAGCGAGTCAGTGAGCGAGGAAGCGGAAGAGCGCCCAATACGCAAACCGCCTCTCCCCGCGCGTTGGCCGATTCATTAATGCAGCTGGCACGACAGGTTTCCCGACTGGAAAGCGGGCAGTGAGCGCAACGCAATTAATGTGAGTTAGCTCACTCATTAGGCACCCCAGGCTTTACACTTTATGCTTCCGGCTCGTATGTTGTGTGGAATTGTGAGCGGATAACAATTTCACACAGGAAACAGCTATGACCATGATTACGCCAAGCTCTAATACGACTCACTATAGGGAAAGCTCGGTACCACGCATGCTGCAGACGCGTTACGTATCGGATCCAGAATTCGTGATGATCTTAAGGCTAGAGTACTAATACGACTCACTATAGGGAGACCACAACGGTTTCCCTCTAGAAATAATTTTGTTTAACTTAAGAAGGAGGAAAAAAAAATGTCTAAAGGTGAAGAATTATTCACTGGTGTTGTCCCAATTTTGGTTGAATTAGATGGTGATGTTAATGGTCACAAATTTTCTGTCTCCGGTGAAGGTGAAGGTGATGCTACTTACGGTAAATTGACCTTAAAATTTATTTGTACTACTGGTAAATTGCCAGTTCCATGGCCAACCTTAGTCACTACTTTAACTTATGGTGTTCAATGTTTTTCTAGATACCCAGATCATATGAAACAACATGACTTTTTCAAGTCTGCCATGCCAGAAGGTTATGTTCAAGAAAGAACTATTTTTTTCAAAGATGACGGTAACTACAAGACCAGAGCTGAAGTCAAGTTTGAAGGTGATACCTTAGTTAATAGAATCGAATTAAAAGGTATTGATTTTAAAGAAGATGGTAACATTTTAGGTCACAAATTGGAATACAACTATAACTCTCACAATGTTTACATCATGGCTGACAAACAAAAGAATGGTATCAAAGTTAACTTCAAAATTAGACACAACATTGAAGATGGTTCTGTTCAATTAGCTGACCATTATCAACAAAATACTCCAATTGGTGATGGTCCAGTCTTGTTACCAGACAACCATTACTTATCCACTCAATCTGCCTTATCCAAAGATCCAAACGAAAAGAGAGACCACATGGTCTTGTTAGAATTTGTTACTGCTGCTGGTATTACCCATGGTATGGATGAATTGTACAAATAATAACGACTCAGGCTGCTACGCCTGTGTACTGGAAAACAAAACCAAAACCCAAAAAACAAAAAACTGAGCCCATTGGTATCGTGGAAGGACTCTATCAAAAAAAAAAAAAAAAAAAAAAAAAAAAAACTAGCATAACCCCTTGGGGCCTCTAAACGGGTCTTGAGGGGTTTTTTGATCTGAATTCGTCGACAAGCTTCTCGAGCCTAGGCTAGCTCTAGACCACACGTGTGGGGGCCCGAGCTCGCGGCCGCTGTATTCTATAGTGTCACCTAAATGGCCGCACAATTCACTGGCCGTCGTTTTACAACGTCGTGACTGGGAAAACCCTGGCGTTACCCAACTTAATCGCCTTGCAGCACATCCCCCTTTCGCCAGCTGGCGTAATAGCGAAGAGGCCCGCACCGATCGCCCTTCCCAACAGTTGCGCAGCCTGAATGGCGAATGGAAATTGTAAGCGTTAATATTTTGTTAAAATTCGCGTTAAATTTTTGTTAAATCAGCTCATTTTTTAACCAATAGGCCGAAATCGGCAAAATCCCTTATAAATCAAAAGAATAGACCGAGATAGGGTTGAGTGTTGTTCCAGTTTGGAACAAGAGTCCACTATTAAAGAACGTGGACTCCAACGTCAAAGGGCGAAAAACCGTCTATCAGGGCGATGGCCCACTACGTGAACCATCACCCTAATCAAGTTTTTTGGGGTCGAGGTGCCGTAAAGCACTAAATCGGAACCCTAAAGGGAGCCCCCGATTTAGAGCTTGACGGGGAAAGCCGGCGAACGTGGCGAGAAAGGAAGGGAAGAAAGCGAAAGGAGCGGGCGCTAGGGCGCTGGCAAGTGTAGCGGTCACGCTGCGCGTAACCACCACACCCGCCGCGCTTAATGCGCCGCTACAGGGCGCGTCAGGTGGCACTTTTCGGGGAAATGTGCGCGGAACCCCTATTTGTTTATTTTTCTAAATACATTCAAATATGTATCCGCTCATGAGACAATAACCCTGATAAATGCTTCAATAATATTGAAAAAGGAAGAGTATGAGTATTCAACATTTCCGTGTCGCCCTTATTCCCTTTTTTGCGGCATTTTGCCTTCCTGTTTTTGCTCACCCAGAAACGCTGGTGAAAGTAAAAGATGCTGAAGATCAGTTGGGTGCACGAGTGGGTTACATCGAACTGGATCTCAACAGCGGTAAGATCCTTGAGAGTTTTCGCCCCGAAGAACGTTTTCCAATGATGAGCACTTTTAAAGTTCTGCTATGTGGCGCGGTATTATCCCGTATTGACGCCGGGCAAGAGCAACTCGGTCGCCGCATACACTATTCTCAGAATGACTTGGTTGAGTACTCACCAGTCACAGAAAAGCATCTTACGGATGGCATGACAGTAAGAGAATTATGCAGTGCTGCCATAACCATGAGTGATAACACTGCGGCCAACTTACTTCTGACAACGATCGGAGGACCGAAGGAGCTAACCGCTTTTTTGCACAACATGGGGGATCATGTAACTCGCCTTGATCGTTGGGAACCGGAGCTGAATGAAGCCATACCAAACGACGAGCGTGACACCACGATGCCTGTAGCAATGGCAACAACGTTGCGCAAACTATTAACTGGCGAACTACTTACTCTAGCTTCCCGGCAACAATTAATAGACTGGATGGAGGCGGATAAAGTTGCAGGACCACTTCTGCGCTCGGCCCTTCCGGCTGGCTGGTTTATTGCTGATAAATCTGGAGCCGGTGAGCGTGGGTCTCGCGGTATCATTGCAGCACTGGGGCCAGATGGTAAGCCCTCCCGTATCGTAGTTATCTACACGACGGGGAGTCAGGCAACTATGGATGAACGAAATAGACAGATCGCTGAGATAGGTGCCTCACTGATTAAGCATTGGTAACTGTCAGACCAAGTTTACTCATATATACTTTAGATTGATTTAAAACTTCATTTTTAATTTAAAAGGATCTAGGTGAAGATCCTTTTTGATAATCTCATGAACAATAAAACTGTCTGCTTACATAAACAGTAATACAAGGGGTGTTATGAGCCATATTCAACGGGAAACGTCTTGCTCTAGGCCGCGATTAAATTCCAACATGGATGCTGATTTATATGGGTATAAATGGGCTCGCGATAATGTCGGGCAATCAGGTGCGACAATCTATCGATTGTATGGGAAGCCCGATGCGCCAGAGTTGTTTCTGAAACATGGCAAAGGTAGCGTTGCCAATGATGTTACAGATGAGATGGTCAGACTAAACTGGCTGACGGAATTTATGCCTCTTCCGACCATCAAGCATTTTATCCGTACTCCTGATGATGCATGGTTACTCACCACTGCGATCCCCGGGAAAACAGCATTCCAGGTATTAGAAGAATATCCTGATTCAGGTGAAAATATTGTTGATGCGCTGGCAGTGTTCCTGCGCCGGTTGCATTCGATTCCTGTTTGTAAT

TGTCCTTTTAACAGCGATCGCGTATTTCGTCTCG

